# Bacterial cohesion predicts spatial distribution in the larval zebrafish intestine

**DOI:** 10.1101/392316

**Authors:** Brandon H. Schlomann, Travis J. Wiles, Elena S. Wall, Karen Guillemin, Raghuveer Parthasarathy

**Affiliations:** Institute of Molecular Biology, University of Oregon, Eugene, Oregon, United States of America; Department of Physics and Materials Science Institute, University of Oregon, Eugene, Oregon, United States of America; Humans and the Microbiome Program, Canadian Institute for Advanced Research, Toronto, Ontario M5G 1Z8, Canada

## Abstract

Are there general biophysical relationships governing the spatial organization of the gut microbiome? Despite growing realization that spatial structure is important for population stability, inter-bacterial competition, and host functions, it is unclear in any animal gut whether such structure is subject to predictive, unifying rules, or if it results from contextual, species-specific behaviors. To explore this, we used light sheet fluorescence microscopy to conduct a high-resolution comparative study of bacterial distribution patterns throughout the entire intestinal volume of live, larval zebrafish. Fluorescently tagged strains of seven bacterial symbionts, representing six different species native to zebrafish, were each separately mono-associated with animals that had been raised initially germ-free. The strains showed large differences in both cohesion—the degree to which they auto-aggregate—and spatial distribution. We uncovered a striking correlation between each strain’s mean position and its cohesion, whether quantified as the fraction of cells existing as planktonic individuals, the average aggregate size, or the total number of aggregates. Moreover, these correlations held within species as well; aggregates of different sizes localized as predicted from the pan-species observations. Together, our findings indicate that bacteria within the zebrafish intestine are subject to generic processes that organize populations by their cohesive properties. The likely drivers of this relationship, peristaltic fluid flow, tubular anatomy, and bacterial growth and aggregation kinetics, are common throughout animals. We therefore suggest that the framework introduced here, of biophysical links between bacterial cohesion and spatial organization, should be useful for directing explorations in other host-microbe systems, formulating detailed models that can quantitatively map onto experimental data, and developing new tools that manipulate cohesion to engineer microbiome function.

## Introduction

Dense and diverse communities of microbes reside in the intestines of humans and other animals. Their large impact on processes ranging from digestion to disease progression [1, 2, 3] motivates a great deal of work aiming to uncover determinants of community composition and function. Because of the size and anatomy of the gut, and because of the remarkable number of microbial species that coexist within it—hundreds to thousands in humans—it is widely believed that spatial organization plays an important role in orchestrating community structure [4, 5]. In support of this, for example, recent studies have shown that distinct groups of bacteria inhabit the lumenal space of the intestine compared to the dense mucus layer lining the epithelium [6] and that distinct taxa are found in different regions along the length of the digestive tract [7]. The drivers of spatial organization are most often considered to be anatomical, as above, or biochemical, for example caused by variation in pH or the concentrations of nutrients, oxygen, or antimicrobial peptides [8].

Here, we suggest and demonstrate that the biophysical character of the microbes themselves, namely the degree to which they are planktonic or aggregated, can be a strong predictor of their populations overall position within the intestine. In macroscopic ecological contexts, such relationships between morphology and spatial distribution are well known. For example, animal body mass is greater in colder regions (Bergmann’s rule), likely due to the scaling of surface driven heat loss with size; phytoplankton aggregation is correlated with position in the water column, due to buoyancy [9]; and seed mass varies robustly with latitude, for reasons that are still unclear [10].

It remains an open question whether gut microbes are governed by broad, pan-species principles linking cellular behavior and large-scale distribution, or whether spatial structure is contingent on context- and species-specific interactions. Investigating this requires high-resolution imaging within live animals in a controlled setting, which has only recently become possible. Uncovering such principles would demonstrate that despite the biochemical complexity of the vertebrate microbiota, general biophysical principles governing the architecture of gut microbial communities may exist.

Our study makes use of larval zebrafish (Fig. 1A, 1B), a model organism of particular utility to investigations of host-microbe interactions due to its anatomical and physiological similarities to other vertebrates, its optical transparency, and its amenability to gnotobiotic techniques for the creation of fish colonized only by particular microbial species [11, 12, 13, 14]. Zebrafish naturally associate with a diverse intestinal microbiome containing hundreds of bacterial species [15, 16] that influence a wide range of host processes [17, 18, 19, 20]. Earlier work on the dynamics of two native zebrafish bacterial symbionts [14] and a human-derived pathogen [13] showed associations between cellular growth mode, specifically whether the bacteria are planktonic or aggregated, and spatial distribution, specifically the location of the population along the length of the intestine, but the robustness and generality of this association remains unexplored.

**Figure 1:**
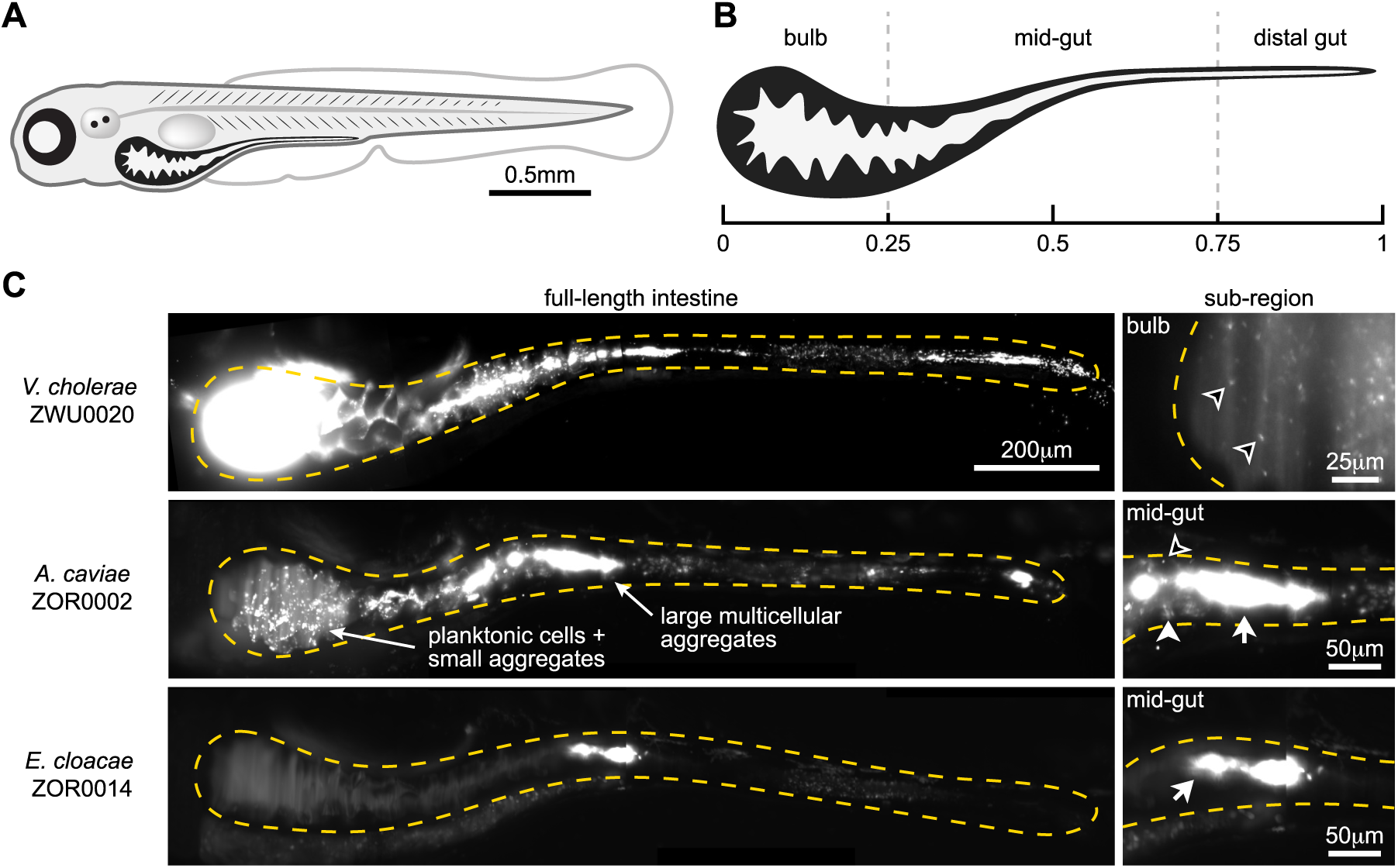
Diversity of bacterial population structures within the zebrafish intestine. (A) Schematic of a 6-day old larval zebrafish. (B) Schematic of a larval zebrafish intestine with the three general anatomical regions and their approximate relative sizes highlighted. (C) Representative images from across the range of observed population structures. Each image is a maximum intensity projection of a full 3D image stack, except for the top right inset, which is a single optical plane. Dashed amber lines trace the approximate boundaries of the intestine in each image. Examples of single cells (open arrowheads), small aggregates (closed arrowheads), and large aggregates (tailed arrowheads) are noted within insets under “sub-region”. See also Supplementary Movies 1-4. Top row: Populations of *V. cholerae* ZWU0020 localize to the anterior bulb and are dominated by highly motile planktonic cells (Supplementary Movie 1). Inset shows *V. cholerae* ZWU0020 cells in a different fish that was colonized with 1:100 mixture of green and red variants. The dilute channel (green) is shown. Middle row: Populations of *A. caviae* ZOR0002 typically contain a range of bacterial aggregate sizes, as indicated by arrows. Inset shows a zoomed-in view of the same intestine. Bottom row: Populations of *E. cloacae* ZOR0014 typically consist of small numbers of large aggregates. Inset shows a zoomed-in view of the same intestine.

### Experimental Design

To investigate this putative relationship, we analyzed seven bacterial strains representing six different species (Table 1). All were isolated from zebrafish intestines, where they are common and abundant [16]. Each species was previously engineered to constitutively express fluorescent proteins [21]. To deduce relationships intrinsic to species morphology and its interaction with the gut environment, and to reduce variation arising from inter-bacterial interactions, we first raised larvae germ-free and then colonized them with individual bacterial strains by inoculation of the aqueous medium (Methods and Supplemental Text). After a 24 hour period of colonization and growth, three-dimensional image stacks were acquired using a custom-built light sheet fluorescence microscope described in detail elsewhere [12]. The images span the entire larval intestine, roughly 200 x 200 x 1000 microns in extent, with single-bacterium resolution. Additional details are provided in Methods.

**Table 1:** Bacterial strains and imaging-derived estimates of mono-association abundances *in vivo*. Abundances were estimated from 3D image stacks using the computational pipeline described in Methods and Supplementary Text

| strain | median abundance |
| --- | --- |
| *Aeromonas veronii* ZOR0001 | $8.4 \times 10^2$ |
| *Aeromonas caviae* ZOR0002 | $1.2 \times 10^3$ |
| *Enterobacter cloacae* ZOR0014 | $3.5 \times 10^3$ |
| *Plesiomonas* ZOR0011 | $4.6 \times 10^3$ |
| *Pseudomonas mendocina* ZWU0006 | $3.5 \times 10^2$ |
| *Vibrio cholerae* ZOR0036 | $1.6 \times 10^3$ |
| *Vibrio cholerae* ZWU0020 | $2.0 \times 10^4$ |

## Results

Imaging multiple fish per strain revealed a broad spectrum of growth modes and bacterial distributions, ranging from the highly planktonic populations of *Vibrio cholerae* ZWU0020 located within the anterior bulb (Fig. 1C, top; Supplemental Movie 1) to the almost entirely aggregated populations of *Enterobacter cloacae* ZOR0014 located within the midgut (Fig. 1C, bottom). Most populations displayed intermediate mixtures of cellular growth modes and spatial distributions, similar to that of *Aeromonas caviae* ZOR0002 (Fig. 1C, middle; Supplemental Fig. 1, Supplemental Movies 2-4). As with observations of *Aeromonas* strains in earlier work [14], bacterial aggregates were dense, compact, and cohesive. The predominant difference in spatial position between species was their location along the longitudinal axis of the intestine. We observed no strains, for example, that localized along the radial axis, lining the gut epithelium.

We computationally identified each individual bacterium and aggregate in each three-dimensional image stack, and also determined the number of cells in each aggregate [12] (see Methods). For each population, we computed the center of mass along the longitudinal axis of the intestine, normalized by the total intestinal length, to represent its spatial distribution. We also enumerated the fraction of the population present as planktonic cells to represent the strains growth mode. Plotting each strain’s planktonic fraction versus its population center shows a clear and striking correlation (Fig. 2A). Linear regression of log_10_-transformed planktonic fraction (log_10_ *f_p_*) against center of mass position (*x_c_*) gives a coefficient of determination of *R*^2^ = 0.91, and best-fit parameters

**Figure 2:**
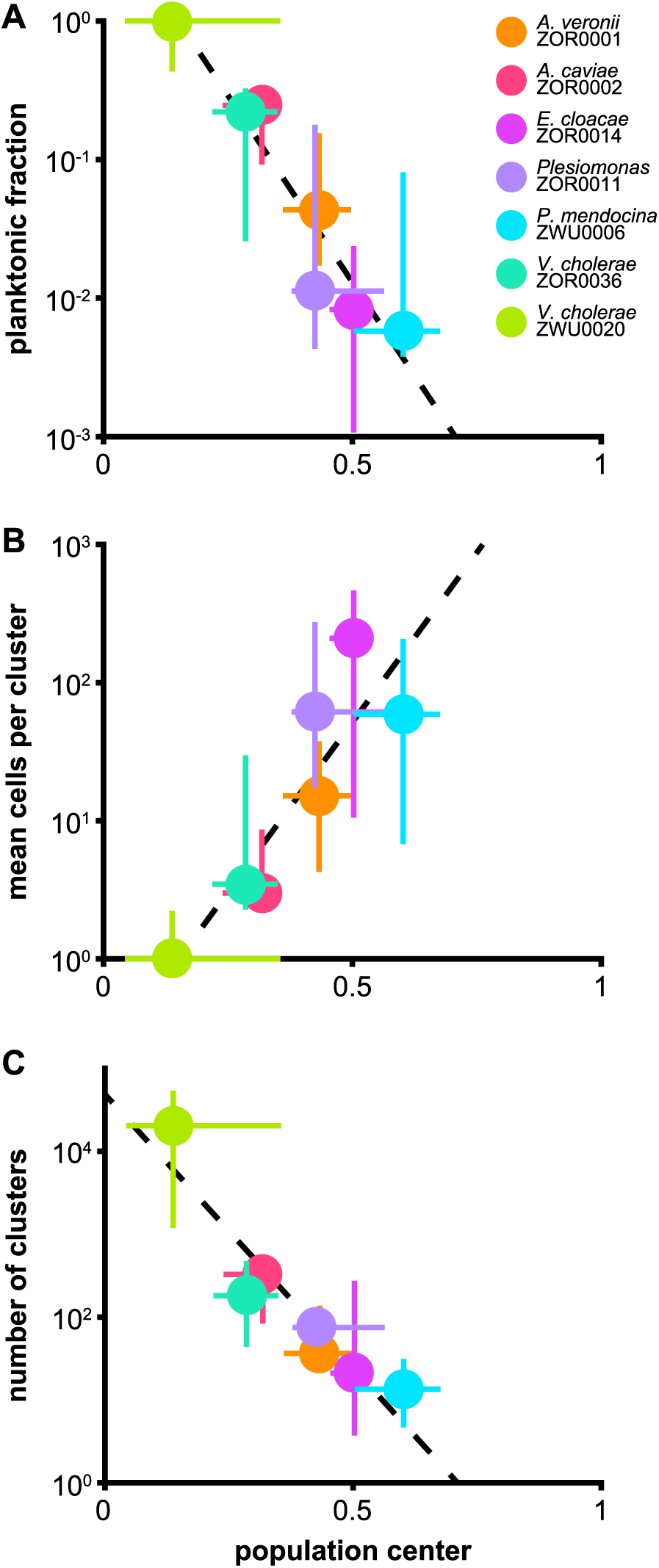
Metrics of cohesion correlate with spatial distribution across bacterial strains. (A) the fraction of the population of each strain existing as single planktonic cells, (B) the average number of cells per cluster, and (C) the total number of clusters plotted against the population center, the center of mass position of each strain normalized by the length of the intestine. For the plots shown in B and C, individual cells are considered clusters of size 1. Circles show median values for each strain, bars show 25% and 75% quartiles. Trendlines were generated from the unweighted linear regression of log_10_-transformed medians.

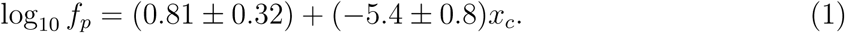

Making use of our image segmentation of bacterial aggregates, we examined the relationship between mean object size and position. Defining a cluster as any group of bacteria (so that an individual bacterium is a cluster of size one), we find a strong correlation between each species’ average cluster size (mean cells per cluster, *C_c_*) and its center of mass (Fig. 2B, *R*^2^= 0.79);

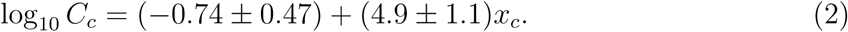

Because *C_c_* is proportional to the total number of cells and inversely proportional to the number of clusters per fish (*n_c_*), the relationship in Fig. 2B could be caused by a dependence on either or both of these factors. However, the total population of each species, save for *V. cholerae* ZWU0020, is roughly similar (Table 1); in contrast, *n_c_* is strongly negatively correlated with position (Fig. 2C, *R*^2^ = 0.88). Regression gives

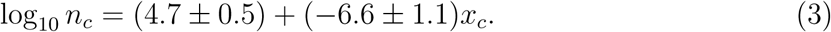

The slope, −6.6 ± 1.1, is close to the negative of the slope of the *C_c_* vs *x_c_* relationship (4.9 ± 1.1), as would be expected if *C_c_* ∼ 1*/n_c_* with the overall population being species-independent. Together, the *C_c_* vs *x_c_* and *n_c_* vs *x_c_* relationships confirm the lack of a global correlation between abundance and location and imply instead that local interactions relate the size and positioning of aggregates.

We next asked if the relationship between aggregation and intestinal distribution we found between strains could be detected within individual strain populations, which would further support its biophysical generality. For this, we considered only clusters of two or more cells because individual cells dominate each dataset (Fig. 2A). For each strain, excluding *V. cholerae* ZWU0020 because it shows almost no aggregation (Fig. 2A), we combined measurements of cluster size and intestinal position from all specimens. We restricted our analysis to the anterior half of the intestine because the distal half rarely contained substantial populations (likely due to frequent intestinal expulsion), limiting our statistical power in that region. Regressing log_10_-transformed sizes of aggregates against their position (Fig. 3, small circles and dashed trendlines), we found a positive correlation between aggregate size and aggregate position for each of the six strains (Table 2). Finding this relationship within strains, as well as across strains, suggests a generic mechanism that spatially segregates bacterial cells based on their cohesive properties, resulting in the localization of small aggregates to the anterior of the intestine and larger aggregates to the posterior.

**Table 2:** Results of within-strain regressions in Fig 3. The regression was of log_10_-transformed cluster sizes (with size ≥ 2) against normalized cluster positions.

| strain | slope value | uncertainty |
| --- | --- | --- |
| *Aeromonas veronii* ZOR0001 | 1.5 | 0.6 |
| *Aeromonas caviae* ZOR0002 | 0.5 | 0.2 |
| *Enterobacter cloacae* ZOR0014 | 1.5 | 0.4 |
| *Plesiomonas* ZOR0011 | 1.4 | 1.5 |
| *Pseudomonas mendocina* ZWU0006 | 1.1 | 1.4 |
| *Vibrio cholerae* ZOR0036 | 0.8 | 0.3 |

**Figure 3:**
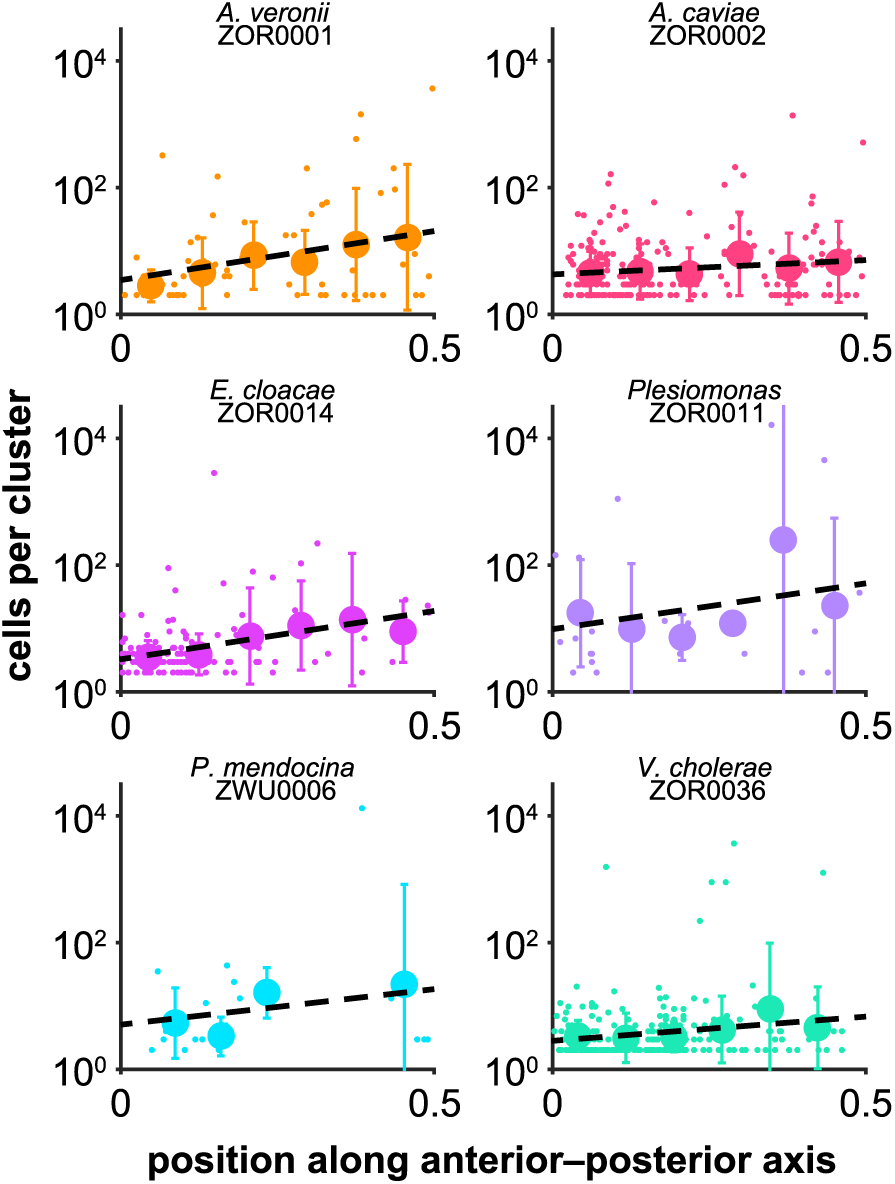
Signatures of a cohesion-distribution relationship can be detected within populations at the strain level. For each strain shown, the size of every cluster with size 2 cells and greater across all samples is plotted against its normalized position along the intestine (small circles). Trendlines depict linear regressions of log_10_-transformed cluster sizes against position (black dashed lines). To better highlight trends, data were binned by position and the mean standard deviation of cluster sizes were overlaid on each plot as large circles and bars.

## Discussion

Harnessing the natural variation displayed by native zebrafish symbionts and the spatial insights made possible by 3D live imaging, we have uncovered a quantitative relationship between bacterial cell behavior and large-scale spatial organization throughout the intestine. We found that across species and strains, the degree to which bacterial populations are aggregated, a biophysical characteristic we term “cohesion”, correlates strongly with their mean position along the intestine. Moreover, looking within strains we were able to detect further signatures of the cohesion-distribution correlation: namely, the size and location of individual aggregates are also correlated. These findings suggest that the relationship between cohesion and spatial structure represents a general principle that manifests across both taxonomic and cellular scales. Intriguingly, the diverse species and strains we examined each have well-defined characteristics, while together they span the range from almost wholly planktonic to almost wholly aggregated, with the corresponding range of intestinal locations. This suggests that either through evolution of particular colonization strategies or behavioral responses to the gut environment, bacteria have the capacity to influence how the intestine shapes their populations.

We posit that the mechanism underlying the cohesion-distribution relationship emerges from the interplay between physical properties of the intestinal environment, especially its shape and peristaltic activity, and the cellular lifestyles of resident bacteria. As in all vertebrates, the larval zebrafish intestine is roughly tubular with a corrugated surface of villi, and transports and mixes contents using coordinated, periodic peristaltic contractions [22]. Earlier work looking solely at *A. veronii* ZOR0001 found aggregated microbes pushed and sporadically ejected by these contractions [14]; such forces more generally affect all aggregated bacteria. Theoretical studies of particle suspensions under low Reynolds number peristaltic flow also show spatial segregation of planktonic and aggregated cells [23]. These observations suggest that it should be possible to construct models that quantitatively match in vivo measurements and that offer predictions relevant for other animals, including humans. The development of such models will be challenging, as they must combine fluid dynamics, anatomy, and the nucleation, growth, and transport properties of bacterial aggregates. Aggregation kinetics are quantifiable from in vivo time-series imaging [12], and ongoing work, from both imaging and modeling, suggests that a robust, pan-species characterization of cluster dynamics is possible.

Even in the absence of such detailed models, however, it is reasonable to believe that the general relationship uncovered here will occur in larger systems, such as the human gut. Peristaltic transport, a tube-like geometry, and bacterial growth are universal features of all animal intestines. Given that Reynolds and Stokes numbers are low in both the zebrafish intestine and the much larger human intestine, we expect that the flow fields and particle transport that result from peristaltic contractions will be similar across scales. This similarity has already allowed quantitative comparisons of microbial compositions driven by pH and flow rates between in vitro fluidic devices and the human microbiome [24]. Therefore, the longitudinal segregation of bacterial clusters by size that we observed here may be a generic consequence of peristaltic activity. Moreover, the finer-scale structure of crypts and folds affords still further possibilities for spatial structuring driven by the associated flow fields and bacterial cohesion. Host anatomy, diet, and biochemical heterogeneity will likely complicate this picture, but we suggest that a general trend connecting bacterial morphology and intestinal position is reasonable to expect and intriguing to search for.

The relationship between cohesion and spatial distribution described here offers a framework for precision microbiome engineering. For example, by manipulating cohesion it may be possible to selectively displace bacterial populations from certain regions of the intestine or to remove them entirely. Reflecting this point, it was recently shown in a murine *Salmonella* vaccine model that antibody-mediated enchaining of bacterial cells led to aggregation and intestinal expulsion [25]. In addition, peristaltic activity can change in response to diet, therapeutic drugs, infection, and a range of chronic diseases. Therefore, elaborating the link between cohesion, spatial structure, and flow may help explain diseases that result from microbial imbalances, and inspire methods for countering such changes in community composition through the targeted alteration of bacterial aggregation.

## Methods

### Bacteria

All bacterial strains used in this study are listed in Table 1. Each strain was previously engineered via Tn7-mediated insertion to constitutively express either dTomato or sfGFP fluorescent reporters from a single chromosomal locus [21]. Archived stocks of bacteria were maintained in 25% glycerol at −80C. Prior to experiments, bacteria were directly inoculated from frozen stocks into 5ml lysogeny broth (LB) media (10g/L NaCl, 5g/L yeast extract, 12g/L tryptone, 1g/L glucose) and grown for 16h (overnight) shaking at 30C.

### Animal care and gnotobiology

All experiments with zebrafish were done in accordance with protocols approved by the University of Oregon Institutional Animal Care and Use Committee and following standard operating procedures [26]. Wild-type (AB x TU strain) zebrafish were derived germ-free and colonized with bacterial strains as previously described [27] with slight modification (Supplemental Text).

### Live imaging

Live imaging of larval zebrafish was conducted using a home-built light sheet fluorescence microscope previously described in detail [12]. The full volume of the intestine (approximately 1200×300×150 microns) is captured in four sub-regions that are registered in software following imaging. An entire intestine sampled with 1-micron steps between planes is imaged in less than 1 minute. All images were taken with an exposure time of 30ms and an excitation laser power of 5mW at 488 nm and 561 nm wavelengths.

### Image analysis

Three-dimensional image stacks were analyzed using a pipeline described in detail in [12], with minor changes (Supplemental Text). The goal of the analysis is to identify the location and size of all bacterial clusters, ranging from individual, planktonic cells to large multicellular aggregates. Small objects are identified using a spot detection algorithm calibrated to over count spots, which are then filtered using a trained classifier (Supplemental Text). Large objects are segmented using a graph-cut algorithm [28], typically seeded with a mask obtained by intensity thresholding. The number of cells per multicellular aggregate is estimated by dividing the total fluorescence intensity of the aggregate by the average intensity of single cells in the same fish host. In cases where single cells are sparse or absent, the average is taken across all single cells for that strain.

### Data

To maximize statistical power, we combined newly acquired data with a recently published image dataset obtained under identical conditions [21]. The recently published data had been subjected to prior analysis to estimate overall bacterial abundances, but was reanalyzed here from scratch using the methods described above and in the Supplemental Text. The combined dataset consisted of N=6 fish per strain, except for *Plesiomonas* ZOR0011, which had N=3 fish. The output of our computational pipeline, a text file containing the size and location of every bacterial cluster, with identifiers for strain, fish, and dataset, is included in Supplemental Data File 1, with details on its format in the Supplemental Text. In addition, a spreadsheet with the cohesion and distribution metrics plotted in Figure 2 is included in Supplemental Data File 2.

## Acknowledgements

Research was supported through the M.J. Murdock Charitable Trust and an award from the Kavli Microbiome Ideas Challenge, a project led by the American Society for Microbiology in partnership with the American Chemical Society and the American Physical Society and supported by The Kavli Foundation. Work was also supported by the National Science Foundation under Award 1427957. Authors received funding from the National Institutes of Health (NIH, http://www.nih.gov/), P50GM09891 to KG and RP, F32AI112094 to TJW, and T32GM007759 to BHS. The funders had no role in study design, data collection and analysis, decision to publish, or preparation of the manuscript.

## Supplemental Text

### Zebrafish gnotobiology details

Wild-type (AB x TU strain) zebrafish were derived germ-free and colonized with bacterial strains as previously described [27] with slight modification. Briefly, fertilized eggs from adult mating pairs were collected and incubated in filter-sterilized embryo media (EM) containing ampicillin (100 *µ*g/ml), gentamicin (10 *µ*g/ml), amphotericin B (250 ng/ml), tetracycline (1 *µ*g/ml), and chloramphenicol (1 *µ*g/ml) for ~ 6h. Embryos were next washed in EM containing 0.1% polyvinylpyrrolidone-iodine followed by EM containing 0.003% sodium hypochlorite. Surface-sterilized embryos were distributed into T25 tissue culture flasks containing 15 ml sterile EM at a density of one embryo per ml and incubated at 28-30^*◦*^ C prior to bacterial colonization. Embryos were sustained on yolk-derived nutrients and not fed during experiments. For bacterial mono-association, bacteria were grown overnight in 5ml LB liquid media with shaking at 30^*◦*^C, and prepared for inoculation by pelleting 1ml of dense culture for 2min at 7,000×g, washing once in sterile EM. Each bacterial strain was individually inoculated into the water column of single flasks containing 4-day old germ-free larval zebrafish at a final density of ~ 10^6^ bacteria/ml. Live imaging of bacterial colonization patterns was assessed 24h later.

### Image analysis details

Three-dimensional fluorescence images of bacteria in the zebrafish intestine were segmented using a computational pipeline described in [12] with minor but relevant changes elaborated here. The goal of the pipeline is to identify the size and location of every bacterial cluster in the intestine, ranging from single, planktonic cells to large multicellular aggregates. The original pipeline was calibrated on images of a single bacterial strain. Applied to the diverse set of strains used in this study, we found that its performance varied by strain and deemed the overall accuracy inadequate for the cluster size measurements we sought. Therefore, we introduced changes that led to greater generality at the expense of a small increase in human input. Ongoing efforts are directed at simultaneously maximizing automation and accuracy.

### Single cell detection

In the pipeline, single cells (small objects) and aggregates (large objects) are identified separately and the results are then merged. Putative single cells are identified using a spot detection algorithm calibrated to over-count cells and are then filtered using a trained classifier. In the original pipeline, a wavelet filtering-based algorithm [29] was used identify putative cells. When optimized, this algorithm typically leads to few false positive. For certain strains, however, we found that it under counted cells. We therefore replaced it with a less sophisticated and faster Difference of Gaussians (DoG) algorithm that led to more false positives but rarely missed true cells.

### Single cell classification

In past work [12], we further discriminated between bacteria and non-bacteria by training Support Vector Machine (SVM) classifiers on subsets of human classified data, with one classifier per strain. In the present study, however, training data was sparse for most strains, limiting the power of any machine learning algorithm. To compensate for this, we trained a single SVM classifier on a pooled data set of all strains, with two fish per strain, containing 7000 objects in total. With 11 features (various intensity and shape metrics) and a Radial Basis Function kernel we reached a maximum out of sample accuracy of 68±2%, as measured by cross validation on 90/10 train-test splits. While this accuracy is quite poor, the pooled classifier along with the DoG spot detection algorithm led to improved overall accuracy of the single cell detection pipeline compared to the original wavelet filtering-based approach, assessed by eye. For comparison, SVM classifiers trained on just a single strain using datasets of up to ∼20,000 objects have an average accuracy of ∼75% [30].

As a final step, coarse human classification was performed on the remaining spots. A recent study [30] on the performance of various machine learning algorithms on this task measured a maximum possible accuracy as the average agreement between several human experts on a single dataset, which was ∼90%. As we followed identical protocols, we estimate the accuracy of our spot detection to be ∼90%.

### Multicellular aggregate detection

In the original pipeline, multicellular aggregates were segmented using a graph-cut approach [28] seeded with an intensity threshold mask. We found that this approach performed poorly on images of dense collections of small and mid-sized aggregates, and on images with both extremely large and small aggregates. Therefore, in such cases we instead seeded the graph-cut algorithm with a gradient threshold mask, obtained by thresholding a Sobel-filtered image.

### Special handling of *Vibrio cholerae* ZWU0020

For *Vibrio cholerae* ZWU0020, the bulk of the population is contained in a dense swarm of motile individuals in the anterior bulb (Fig 1C, top; Supplementary Movie 1), obscuring detection by our algorithm designed to identify single cells. To enumerate this population, the swarm is identified as a large cluster, normalized by average single cell fluorescence intensity, and then counted as a collection of single cells. The location of each of these cells is taken to be the center of mass of the whole swarm. *Vibrio cholerae* ZWU0020 is the only strain for which this approach is required.

### Details of Supplemental Data File 1

Supplemental Data File 1 is a text file containing the size and normalized location of every bacterial cluster identified in this study. Each row of the file corresponds to a cluster. The six columns are, from left to right: normalized location along length of the intestine, (unrounded) number of cells per cluster, strain identifier, dataset identifier, fish identifier, cluster identifier.

For multicellular aggregates, location corresponds to the center of mass of the aggregate. Strain identifier is a number 1-7 denoting the bacterial strain, with numbers corresponding to *Aeromonas veronii* ZOR0001, *Aeromonas caviae* ZOR0002, *Enterobacter cloacae* ZOR0014, *Plesiomonas* ZOR0011, *Pseudomonas mendocina* ZWU0006, *Vibrio cholerae* ZWU0020, and *Vibrio cholerae* ZOR0036 respectively. The dataset identifier is equal to 1 if the cluster came from the previously published dataset [21], and is equal to 2 if it came from the dataset generated specifically for this study. The fish identifier labels the fish host the cluster belongs to. The cluster identifier is a unique number assigned to each cluster within a fish.

---

**Figure S1:**
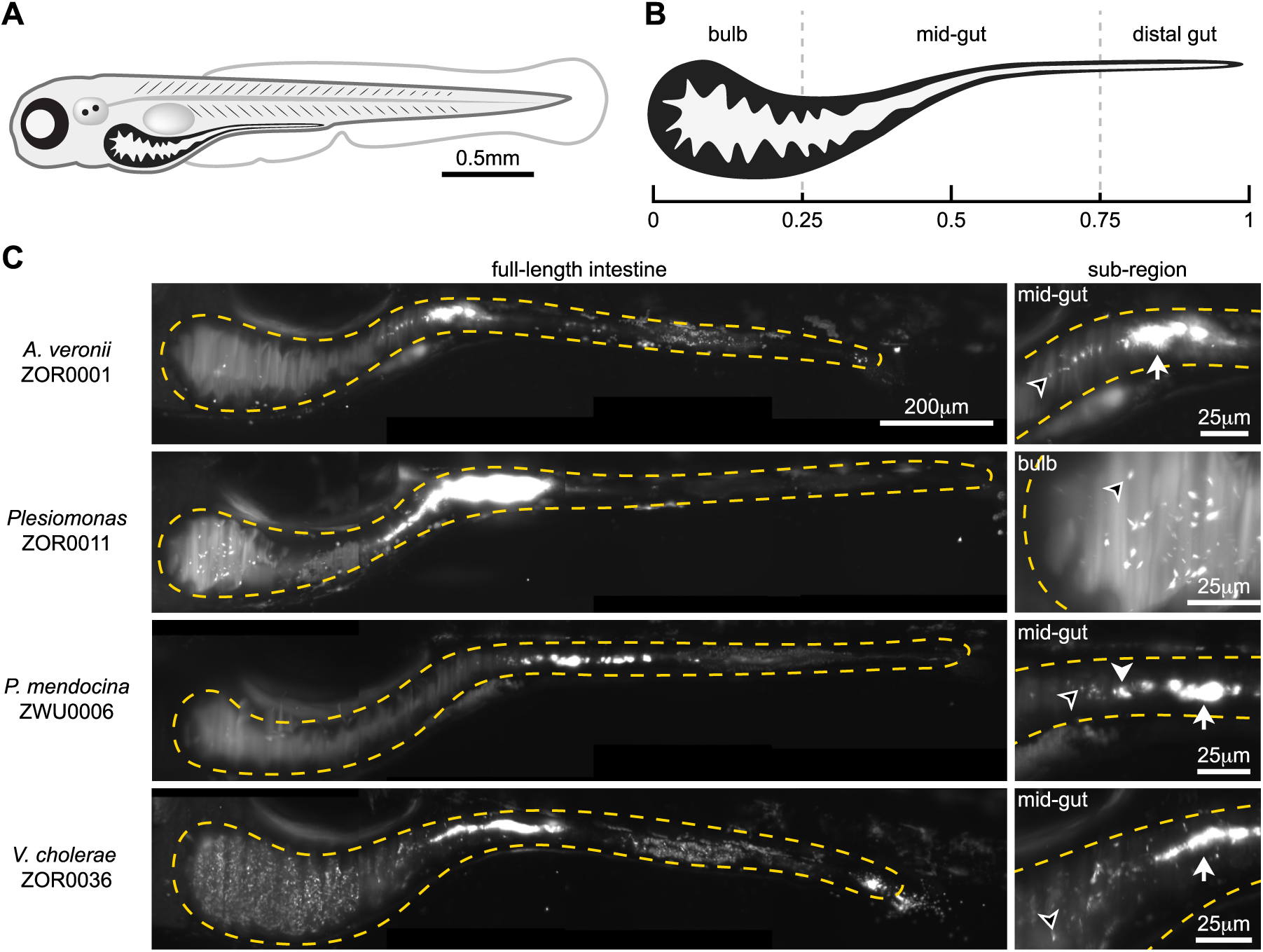
Represetative examples of additional bacterial strains. (A) Schematic of a 6-day old larval zebrafish. (B) Schematic of a larval zebrafish intestine with the three general anatomical regions and their approximate relative sizes highlighted. (C) Representative images from four additional bacterial strains not pictured in the main text. Each image is a maximum intensity projection of a full 3D image stack. Dashed amber lines trace the approximate boundaries of the intestine in each image. Examples of single cells (open arrowheads), small aggregates (closed arrowheads), and large aggregates (tailed arrowheads) are noted within insets under “sub-region”. Each inset is a zoomed in view of the same intestine. First row: Populations of *A. veronii* ZOR0001 localize predominantly to the midgut and feature a mix of planktonic cells and clumps. Second row: Populations of *Plesiomonas* ZOR0011 typically contain a mix of planktonic cells localized in the anterior bulb and aggregates in the midgut. Third row: Populations of *P. mendocina* ZWU0006 typically are dominated by a small number of large aggregates, but planktonic cells are also observed. Fourth row: populations of *V. cholerae* ZOR0036 often contain large numbers of planktonic cells present the anterior bulb, along with more moderately sized aggregates in the midgut.

## Supplemental Movies

### Supplemental Movie 1

Real time movies of two differently tagged variants of *V. cholerae* ZWU0020, inoculated at a 1:100 ratio (GFP:dTomato), in a live, 6-day old zebrafish intestine. Left panel shows a dense swarm of the abundant and highly motile dTomato population. Right panel shows the minor GFP subpopulation, where single cells are more discernible. Movies were captured sequentially, and are from a single optical plane positioned in the anterior bulb. Scale bar = 25 microns.

### Supplemental Movie 2

Animated slices of a z-stack depicting *V. cholerae* ZOR0036 in the anterior bulb and midgut of a 5-day old zebrafish. Beginning at 30 microns in depth, the structure of the intestinal folds and the intestine boundary become visible. Bright puncta within this boundary are bacterial cells. The motion of single cells reflects swimming motility during 3D image acquisition. Vertical stripes in the lumen are the result of shadows cast by pigmented cells on the skin. At 85 microns in depth, strong autofluorescence from the yolk appears in the lower-right corner of the movie. At 125 microns in depth, a large, multicellular aggregate appears in the upper-right corner of the movie, corresponding to the beginning of the midgut. Scale bar = 25 microns.

### Supplemental Movie 3

Animated slices of a z-stack depicting *A. caviae* ZOR0002 in the anterior bulb and midgut of a 5-day old zebrafish. Planktonic cells, small aggregates, and large aggregates are present. In contrast to *V. cholerae* ZOR0036 (Supplemental Movie 2), planktonic cells are not motile in vivo. Vertical stripes in the lumen are the result of shadows cast by pigmented cells on the skin. Scale bar = 25 microns.

### Supplemental Movie 4

3D rendering of *P. mendocina* ZWU0006 in the midgut of a 5-day old zebrafish. A single, large aggregate dominates the field of view. The bacterial aggregate is shown in bright white, whereas autofluorescent intestinal mucus marks the lumenal space. Finer structure is visible, with distinct regions resembling smaller aggregates that appear to have been packed together. Scale bar = 25 microns.

